# Toward sustainable jet fuels: bioconversion of cellulose into isoprenoid biojet candidates using rumen bacteria and non-conventional yeast

**DOI:** 10.1101/2022.07.15.500214

**Authors:** Laura E. Walls, Peter Otoupal, Rodrigo Ledesma-Amaro, Sharon B. Velasquez-Orta, John M. Gladden, Leonardo Rios-Solis

## Abstract

In this study, organic acids were demonstrated as a promising carbon source for bisabolene production by the non-conventional yeast, *Rhodosporidium toruloides*, at microscale with a maximum titre of 1055 ± 7 mg/L. A 125-fold scale-up of the optimal process, enhanced bisabolene titres 2.5-fold to 2606 mg/L. Implementation of a pH controlled organic acid feeding strategy at this scale lead to a further threefold improvement in bisabolene titre to 7758 mg/L, the highest reported microbial titre. Finally, a proof-of-concept sequential bioreactor approach was investigated. Firstly, the cellulolytic bacterium *Ruminococcus flavefaciens* was employed to ferment cellulose, yielding 4.2 g/L of organic acids. *R. toruloides* was subsequently cultivated in the resulting supernatant, producing 318 ± 22 mg/L of bisabolene. This highlights the feasibility of a sequential bioprocess for the bioconversion of cellulose, into biojet fuel candidates. Future work will focus on enhancing organic acid yields and the use of real lignocellulosic feedstocks to further enhance bisabolene production.

**Graphical Abstract:** 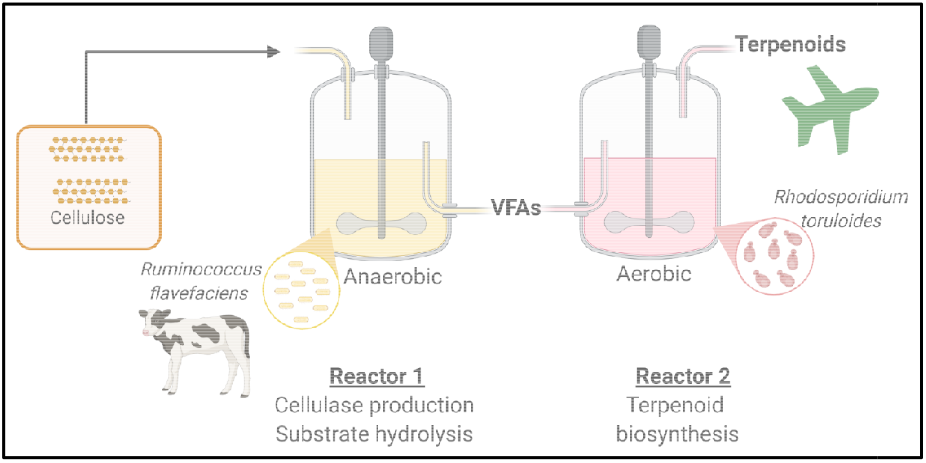

## 1. Introduction

As the fastest mode of transport, the aircraft is a major driver for globalisation and economic growth. However, sustainable development is a major ongoing challenge in the sector. Although the introduction of low emission electric vehicles has been demonstrated as a viable alternative to petroleum based fuels for powering small road vehicles, state-of-the-art lithium ion batteries have an energy density of around 250 Wh/kg(KOKAM, 2019), just 2% of that of the aircraft fuel, Jet A. This capacity is simply insufficient for powering larger vehicles such as those required for aviation, shipping and heavy goods vehicles and as a result reliance on liquid fuels is likely to continue for some time (The Royal Academy of Engineering, 2017). The development of alternative, advanced liquid biofuels with properties similar to petroleum based fuels is therefore crucial meeting the ambitious renewable energy targets recently introduced across Europe (The European Parliament and of the Council of the European Union, 2018; Walls and Rios-Solis, 2020) and the United States (The White House, 2021).

Cellulose, as the most abundant organic material on earth (Carere et al., 2008), holds great promise as a sustainable feedstock for the production of such advanced biofuels. However, the complex crystalline structure thwarts its metabolism by most conventional microbial hosts. As a result, a series of pre-treatment, cellulase production and hydrolysis steps are typically required prior to fermentation of the resulting monomeric sugars (Torres-Sebastián et al., 2021). As traditional approaches involve the use of separate unit operations for each step, economic feasibility is limited for the production of low value chemicals such as biofuels (Navarrete et al., 2020). In addition, 30-50 mg of commercial enzymes are required to hydrolyse each gram of crystalline cellulose (Ali et al., 2016), further increasing operational costs. Process intensification is an innovative engineering approach, which aims to dramatically improve efficiency through considerable reductions in plant volume (ideally 100 to 1000-fold) (Boodhoo and Harvey, 2013). In the context of lignocellulosic biofuel production, process intensification has been achieved through simultaneous saccharification and fermentation (SSF) approaches (Olofsson et al., 2008), in which the enzymatic feedstock degradation and biofuel production steps are combined in a single unit. More recently, a novel one-pot process for the pretreatment, saccharification and bioconversion of sorghum biomass into bisabolene using *R. toruloides* (Sundstrom et al., 2018). Whilst these intensified processes demonstrate considerable progress, the high cost of the required commercial cellulase enzymes still hinders economic feasibility.

Consolidated bioprocessing (CBP) is an innovative approach, which has gained substantial interest due to its potential to simplify and reduce the costs associated with cellulosic biofuel production. In a CBP cellulose production, saccharification and fermentation all take place inside a single unit operation. Ruminants are an excellent example of a natural consolidated bioprocess and can be considered the “largest commercial fermentation process” (2 × 10^11^ L) (Seshadri et al., 2018; Weimer, 1992; Weimer et al., 2009) Ruminants are capable of effective bioconversion of cellulose-rich biomass into valuable products (i.e. meat and milk) within a single process unit (See Supplementary Material).

Within the rumen, a consortium of bacteria, protozoa, archaea and fungi efficiently break down lignocellulosic biomass. The predominant products are volatile fatty acids, which are metabolised by the ruminant, along with CO_2_ and methane. As a result, the rumen is gaining interest as a source of CBP hosts. *Ruminococcus flavefaciens*, for example, is one of the most abundant cellulolytic bacteria in bovine and equine rumen (Julliand et al., 1999; Ozbayram et al., 2018) and is capable of efficient cellulose and hemicellulose fermentation. The predominant products of cellulose fermentation by the species are succinic and acetic acids (Latham and Wolin, 1977). Although, organic acids such as acetic acid are not suitable for advanced liquid biofuel applications (Zuroff et al., 2013), they have been demonstrated as a potential carbon source for oleaginous yeasts (Huang et al., 2016; Liu et al., 2017). The oleaginous yeast, *Rhodosporidium toruloides* (also known as *Rhodotorula toruloides*), for example, was successfully cultivated with 4-20 % acetic acid as the sole carbon source, for the production of lipids (Huang et al., 2016). *R. toruloides* has also gained recent interest as a model host for the production of advanced isoprenoid derived biofuel compounds such as bisabolene, epi-isozizaene and prespatane (Geiselman et al., 2020; Kirby et al., 2021). With properties synonymous with petroleum derived fuels, such compounds hold great promise as “drop-in” replacement jet fuels. As a carotogenic yeast, flux through the mevalonate pathway is considerably higher in these species compared to the traditionally employed yeast, *Saccharomyces cerevisiae*. As a result, through integration of bisabolene synthase genes alone, bisabolene titres of up to 2.2 g/L could be achieved (Kirby et al., 2021). In comparison, extensive mevalonate pathway engineering was required to facilitate the maximum reported titre of 0.99 g/L in *S. cerevisiae* (Peralta-Yahya et al., 2011). In addition, *R. toruloides* is more suited to lignocellulosic hydrolysates as it is capable of simultaneous xylose and glucose metabolism (Zhuang et al., 2019) and has a much greater tolerance for inhibitory compounds such as vanillin and furufal (Yaegashi et al., 2017), which are often generated during the pretreatment step.

A bottleneck in CBP approaches, however, is the stark differences in cultivation requirements of each step (Navarrete et al., 2020; Walls and Rios-Solis, 2020). Most rumen bacteria such as *R. flavefaciens* are obligate anaerobes and grow well at around 37 °C. Most oleaginous yeasts like *R. toruloides*, on the other hand are obligate aerobes and are typically cultivated at a lower temperature of around 30 °C. One potential solution is a sequential bioreactor approach (Navarrete et al., 2020; Walls and Rios-Solis, 2020). To reduce the metabolic burden associated with the expression of the heterologous pathways required for feedstock degradation and product biosynthesis, two microorganisms can be employed, each engineered for optimal performance of one step (Navarrete et al., 2020). Co-cultivation of the two microorganisms in the same reactor may result in competition or necessitate the use of suboptimal operational conditions due to the different requirements of the two species. A sequential bioreactor approach, in which cellulose degradation occurs in the first reactor, followed by bioconversion of the resulting organic acids into the biofuel of interest in a second reactor could effectively overcome these critical challenges.

In this study a proof-of-concept sequential bioprocess was developed for the bioconversion of cellulose to bisabolene using rumen bacteria and non-conventional yeast. The feasibility of succinic and acetic acids as a carbon source for bisabolene production by engineered *R. toruloides* cell factories was assessed in detail at micro-(∼2 mL) and mini-bioreactor (250-500 mL) scale. The ability of *R. flavefaciens* to ferment microcrystalline cellulose under strict anaerobic conditions was also evaluated. Finally, the two steps were combined to test the proposed sequential bioprocess.

## 2. Materials and methods

### 2.1. Microbial strains

The yeast strains used in this study were originally derived from *R. toruloides IFO0880*, mating type A2. Strain GB2 (IFO0880*/*P_*GAPDH*_*-*BIS*-*T_*NOS*_, P_*ANT*_*-*BIS-T_*NOS*_) was engineered for the production of bisabolene through the integration of ten copies of bisabolene synthase into IFO0880 under the control of the native promoter P_GAPDH_ and six copies under the control of P_ANT,_ as described in detail previously (Kirby et al., 2021). Strain GB2.485 (GB2/P_*ANT*_*-* SpHMGR-T_*NOS*__P_*SKP1*_*-*McMK-T_*SKP1*__P_*DUF*_-SpPMK-T_*DUF*_) was then constructed with the aim of improving bisabolene production by GB2 through targeted overexpression of key mevalonate pathway genes, HMGR, MK and PMK in GB2 (Kirby et al., 2021). Further details on strains GB2 and GB2.485 can be found within the Agile BioFoundry Strain Registry (http://public-registry.agilebiofoundry.org), registry IDs ABFPUB_000311 and ABFPUB_000319, respectively. Strain EIZS2 (IFO0880/P_GAPDH_-RtEIZS-NAT^R^) was engineered for the production of epi-isozizaene through the integration of an epi-isozizaene synthase from *Streptomyces coelicolor* codon optimised for expression by *R. toruloides*, under the control of the native GADPH promoter into IFO0880 as described previously (Geiselman et al., 2020). Construction of the prespatane producing strain, PPS5 (IFO0880/P_*ANT*_*-*RtPPS*-*P_*TEF1*_*-*RtPPS-NAT^*R*^/P_*TEF1*_*-*RtPPS-HYG^R^), was achieved through integration of a prespatane synthase gene from *Laurencia pacifica*, codon optimised for expression by *R. toruloides* under the control of native promoters ANT and TEF1 as described in detail previously (Geiselman et al., 2020). Further information on EIZS2 and PPS5 can be found in the Joint BioEnergy Institute Strain Registry (https://public-registry.jbei.org) using registry IDs JPUB_013532 and JPUB_013541, respectively. The biosynthetic pathways for the production of each of the isoprenoid fuel candidates are summarised in Supplementary Material.

The cellulolytic bacteria *Ruminococcus flavefaciens* strain 17, originally derived from bovine rumen, was obtained as a freeze-dried pellet within a double glass ampoule from the German Collection of Microorganisms and Cell Cultures (DSMZ).

### 2.2. Preliminary high-throughput design of experiments guided screening

To assess the feasibility of succinic and acetic acids as carbon sources for the production of bisabolene by *R. toruloides*, a definitive screening design was created using JMP Pro 14 statistical software. The effect of six factors on bisabolene accumulation was considered (See Supplementary Material). Each of the 14 factor combinations included in the experimental design were tested in triplicate in 24-deep-well plates (Axygen, USA). As execution of the design necessitated the use of two 24-well plates across two separate runs, blocking was used to account for any experimental error that may arise as a result. Treatments evaluated in the first plate were assigned block 1 and those in the second were assigned block 2 (Table 1). Inocula preparation was achieved by transferring single colonies of *R. toruloides* strain GB2.485 to 5 mL of rich YPD medium (1% yeast extract; 2% peptone; and 2% glucose) and incubating at 30 °C and 200 rpm overnight. Aliquots of the inocula were diluted with appropriate volumes of each of the media combinations (Table 1) to give a 1.6 mL culture with the desired initial OD_600_. A 400-µL dodecane overlay was added for *in situ* extraction of the bisabolene product. The resulting cultures were incubated at 30 °C and 400 rpm for 96 hours using an Eppendorf ThermoMixer® C thermal mixer. At the end of the cultivation, the contents of each well were centrifuged and the organic layer extracted for analysis of bisabolene production via gas chromatography-mass spectrometry (GC-MS). Biomass accumulation was also evaluated at the end of each cultivation using a Nanodrop 2000c spectrophotometer (Thermofisher Scientific, UK).

**Table 1:**
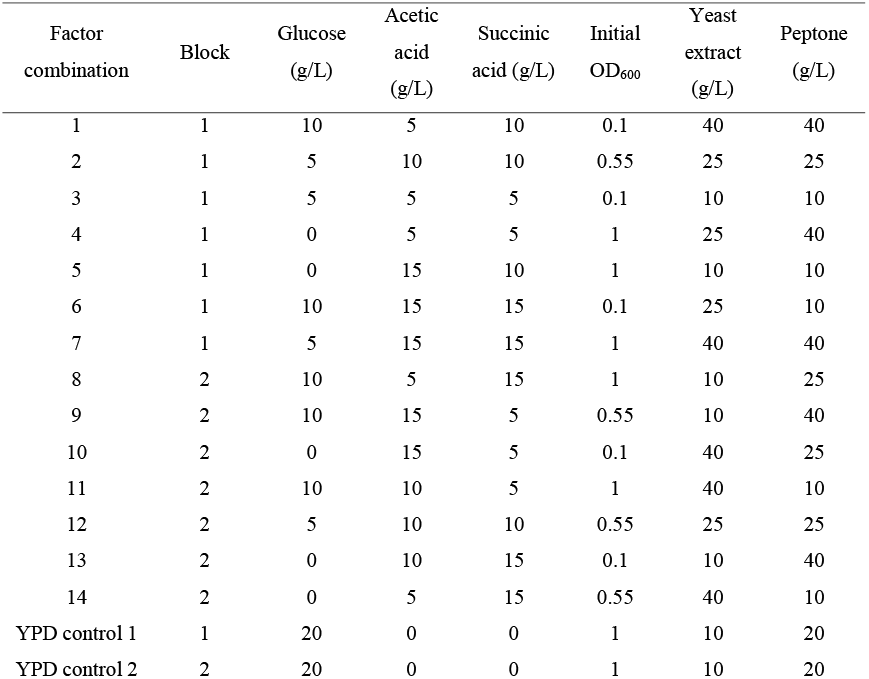
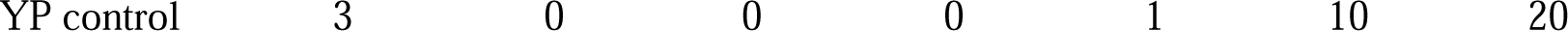
Screening design for high-throughput microscale screening.

### 2.3. Cultivation of Ruminococcus flavefaciens

#### 2.3.1. Cultivation media

*Ruminococcus albus* medium (Medium 436, DSMZ) with minor modifications was used for preliminary cultivations of *Ruminococcus flavefaciens*, containing the following per 100 mL: Peptone 0.5 g; Yeast extract 0.2 g; Glucose 0.3 g/L; Cellobiose 0.2 g; Mineral solution 1 (K_2_HPO_4_ 0.6 %) 7.5 mL; Mineral solution 2 (KH_2_PO_4_ 0.6 %; (NH_4_)_2_SO_4_ 2 %; NaCl 1.2 %; MgSO_4_ X 7 H_2_O 0.25 %; CaCl_2_ X 7 H_2_O 0.16 %) 7.5 mL; Resazurin 0.1 mg; Na_2_CO_3_ 0.4 g; Isobutyric acid 10 µL; Isovaleric acid 10 µL; 2-Methylbutyric acid 10 µL; Cysteine-HCl X H_2_O 50 mg. Once growth was established on Medium 436 *R. flavefaciens* was adapted to growth on microcrystalline cellulose (Avicel PH-101, 50 μm particle size, Sigma-Aldrich, UK) as the main carbon source, firstly the glucose was removed and replaced with 1 g of cellulose (1 % *(w/v)*). In subsequent cultivations the cellobiose was also excluded from the modified medium 436, yielding cellulose medium. For sequential bioprocess cultivations bacteria-yeast media was used, which contained all the elements of the cellulose medium except the concentrations of yeast extract and peptone were increased to 40 and 10 g/L to reflect the optimal concentrations elucidated for *R. toruloides*.

#### 2.3.2. Cultivation conditions

Cultivations of *R. flavefaciens* were performed under strict anaerobic conditions in Hungate type anaerobic cultivation tubes (Sciquip, UK), following a modified Hungate technique (HUNGATE, 1950). Briefly, the cultivation media was boiled under CO_2_ to eliminate oxygen and distributed in 5-7 mL aliquots into Hungate tubes. Each tube was sealed with a butyl rubber stopper and the media was simultaneously heated and bubbled with CO_2_ via a hypodermic needle until elimination of oxygen was confirmed visually due to decolourisation of the resazurin dye. The resulting sealed anoxic tubes were then sterilised via autoclaving prior to use. For cultivation, 0.5 mL of a previous *R. flavefaciens* culture was transferred to each Hungate tube containing anoxic sterile media using a 1 mL syringe and hypodermic needle. Prior to transfer, the syringe was flushed thoroughly with sterile CO_2_ to prevent contamination with oxygen. Following inoculation, the tubes were incubated at 37 °C either with shaking (150-200 rpm) or without shaking as indicated.

#### 2.3.3. Acid quantification

Fermentation of cellulose could be confirmed visually through generation of the characteristic yellow pigment associated with cellulase production in *R. flavefaciens*. Quantification of the major fermentation products, succinic acid and acetic acid, was achieved via enzymatic assay using Succinic Acid Assay Kit (K-SUCC, Megazyme) and Acetic Acid Assay kits (K-ACET, Megazyme), respectively. Prior to analysis, samples were clarified to prevent interference. Briefly, a 100 µL sample was diluted with 700 µL of deionised water before adding 50 µL Carrez solution I (K_3_Fe(CN)_6,_ 3.6 % *(w/v)*), 50 µL of Carrez solution II (ZnSO_4_. 7H_2_O, 7.2 %*(w/v)*) and 100 µL of 100 mM NaOH. Samples were mixed after each addition and the resulting mixture was centrifuged to precipitate contaminants. In both cases, the enzymatic assays were conducted in 96-well plates (Greiner Bio-One) according to the manufacturer’s instructions and absorbance at 340 nm was measured using a CLARIOstar microplate reader (BMG-LABTECH).

### 2.4. Minibioreactor cultivations

#### 2.4.1. Batch validation cultivation

Cultivations were conducted in MiniBio500 bioreactors (Applikon Biotechnology) with a working volume of 250 mL. Pre-inoculum cultures were prepared by transferring from a single GB2.485 colony to 5 mL of YPD and incubating at 30 C and 200 rpm for 8Chours. A 1 mL aliquot of the resulting culture was subsequently used to inoculate a secondary 10 mL YPD culture, which was incubated overnight. An aliquot of the resulting culture was diluted with the optimised acid medium to give a 200 mL culture with an initial OD_600_□= □0.5.

To prevent excess foam production polypropylene glycol P2000 (Alfa Aesar) was added, and a Rushton turbine was placed at the medium-air interface. A 20 % dodecane overlay was added for *in situ* extraction of the isoprenoid products. Temperature, DO, and pH were measured online. The adaptive my-control system (Applikon Biotechnology) was used to control process parameters in the MiniBio 500 bioreactors. A set point of 30 % was implemented for DO and the culture temperature was maintained at 30 C. The pH was maintained around six through the automatic addition of 3 M NaOH or 2 M H_2_SO_4_. Biomass was measured as optical density at 600 nm via offline manual sampling twice daily using a Nanodrop 2000c spectrophotometer (Thermo Fisher Scientific). Samples were also taken twice daily for bisabolene and organic acid quantification via GC-MS and enzymatic assay as per Section 2.4 and Section 2.3.3, respectively.

#### 2.4.2. Feasibility of pH-controlled fed-batch

In the pH-controlled fed-batch cultivation the conditions were as described in Section 2.4.1, however, pH control was achieved using a 5X concentrated organic acid solution (Succinic acid 47.5 g/L, acetic acid 25 g/L) instead of 3 M NaOH and 2 M H_2_SO_4_. The pH was maintained above six through automatic addition of the 5X concentrated acid solution.

### 2.5. Metabolite identification and quantification

Production of bisabolene, epi-isozizaene and prespatane by the engineered *R. toruloides* strains was achieved via gas chromatography – mass spectrometry (GC-MS). Prior to analysis culture samples were centrifuged at 5000 rpm for 5 minutes to separate the aqueous and organic phases. Aliquots of the dodecane overlay containing the respective isoprenoid product were subsequently diluted with dodecane, microplate and initial miniBio 500 samples were diluted fourfold, this dilution was increased up to 20-fold for the miniBio 500 samples to ensure concentrations were within the range of the calibration standards (0 – 1000 mg/L). A 1-μl sample of the resulting mixture was then injected into a TRACE™ 1300 Gas Chromatograph (Thermo Fisher Scientific) coupled to an ISQ LT single quadrupole mass spectrometer (Thermo Fisher Scientific). Chromatographic separation was achieved using a Trace Gold TG-5MS (30 m x 0.25 mm x 0.25 µm) gas chromatography column using a previously described method (Kirby et al., 2021). To confirm production and quantify bisabolene an organic standard solution was used (1000 mg/L, SPEX Certiprep). Production of epi-isozizaene and prespatane was estimated relative to standard concentrations of bisabolene.

## 3. Results and discussion

### 3.1. Feasibility of bisabolene production from organic acids

#### 3.1.1. Design of experiments guided preliminary screening

To investigate the feasibility of organic acids as a carbon source for bisabolene production by an engineered *R. toruloides* strain GB2.485, a design-of-experiments-guided high-throughput screening study was conducted. A definitive screening design (DSD) was selected, as this type of screening design was found to be extremely efficient and informative for the optimisation of isoprenoid production by *Saccharomyces cerevisiae* (Walls et al., 2022, 2021b) The initial concentrations of succinic and acetic acids were deemed critical factors in the study. Previous studies have demonstrated the suitability of acetic acid as a carbon source for lipid (Huang et al., 2016; Robles-Iglesias et al., 2021) and bisabolene production in the species (Magurudeniya et al., 2021; Sundstrom et al., 2018). However, previous studies found that at concentrations above ∼16 g/L acetic acid had a significant inhibitory effect resulting in an extended lag phase and restricted growth (Robles-Iglesias et al., 2021). As tolerance to succinic acid and the suitability of these organic acids for heterologous isoprenoid biosynthesis have been less extensively characterised, these were a focus of the preliminary characterisation study. It was hypothesised that supplementation with a more preferred carbon source may be necessary to generate sufficient biomass for bisabolene synthesis. Glucose supplementation was therefore also included as a factor. Previous studies using *Saccharomyces cerevisiae* microbial cell factories engineered for production of isoprenoid precursors to the chemotherapy drug, Taxol, revealed that the initial biomass concentration had a significant effect on isoprenoid accumulation (Walls et al., 2021a). The initial OD_600_ of the culture was therefore also included as a factor in the design. Finally, the initial concentrations of common complex media components, yeast extract and peptone were considered. The effect of peptone concentration was of particular interest as previous studies have found that carotenoid production is enhanced under nitrogen limited conditions (Zheng et al., 2021). To evaluate the effect of the six factors of interest a DSD was created using JMP Pro 14 statistical software (Table 1). The results of the microscale screening study are summarised in Figure 1.

**Figure 1:**
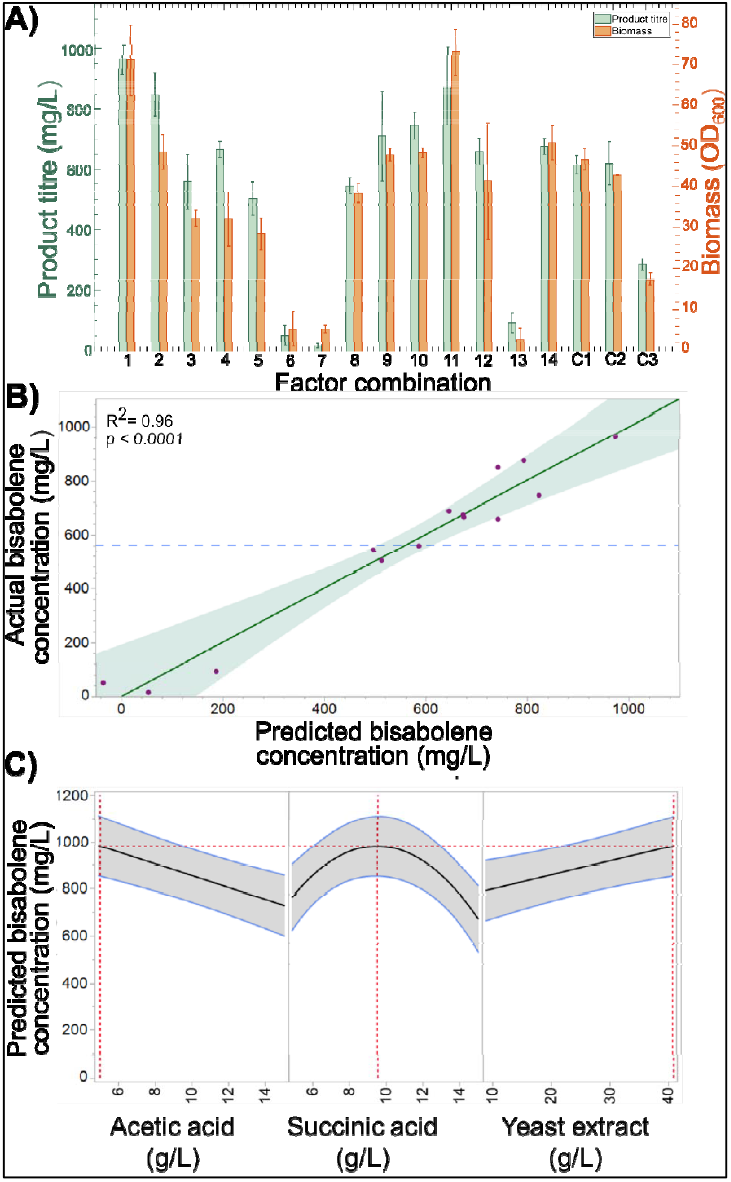
High-throughput design-of-experiments-guided screening summary. A) Biomass and bisabolene accumulation evaluated at the end of 96 hour GB2.485 cultivations performed using each of the 14 media combinations (See Supplementary Material). Control cultivations (C1 and C2) were performed in each of the two 24-deep-well plates using standard YPD media with an initial OD_600_ = 1, an additional control (C3) was performed using YP media alone. Values are mean ± standard deviation for triplicate cultivations. B) Results of the statistical model fitted using JMP Pro 14 statistical software. Experimental bisabolene titres were plotted against those predicted by the statistical model and reliability of the model estimated. Blue dashed line corresponds to mean, solid green line corresponds to fit. C) Optimal setting for the significant main effects as predicted using the prediction profiler and

Of the 14 medium combinations investigated, factor combinations 1, 2 and 11 gave rise to the highest bisabolene titres of 964 ± 47, 849 ± 74 and 874 ± 129 mg/L, respectively. Interestingly, factor combinations 4, 10 and 14 resulted in comparable or slightly higher bisabolene titres to the control YPD cultivations despite using organic acids as the sole carbon source. As the yeast extract and peptone provided an additional source of carbon, a further control cultivation (C3) was performed in which no additional carbon source was added. Bisabolene production was detected with a final titre of 285 ± 19, despite the lack of additional carbon source. To determine which factors had a significant effect on bisabolene production, a statistical model was fitted using the optimised Fit Definitive Screening Design algorithm (Jones and Nachtsheim, 2017) in JMP. Terms with p-values greater than 0.05 were deemed insignificant and therefore manually removed from the model. The final model identified three significant main effects, succinic acid (*p* = 2.0 × 10^−5^), acetic acid (*p* = 3.9 × 10^−4^) and yeast extract (*p* = 6.6 × 10^−3^) concentrations. In addition, an interaction effect was detected between succinic acid and acetic acid concentrations (*p* = 2.2 × 10^−4^). This was expected as previous studies have shown that above organic acid concentrations of around 15-20 g/L, growth is inhibited (Huang et al., 2016; Robles-Iglesias et al., 2021). The effect of succinic acid concentration was also found to be non-linear (p = 6.6 × 10^−4^). The bisabolene titres predicted by the statistical model were plotted against the experimental data using JMP as summarised in Figure 1B. The model achieved an R^2^ of 0.96 and most of the data points were contained within the 95 % confidence interval, indicating a good predictive capability. To determine the optimal settings for each of the factors with a significant effect on the response, the prediction profiler function was used (Figure 1A). According to the statistical model, initial succinic acid, acetic acid and yeast extract concentrations of 9.5, 5 and 40 g/L, respectively, would give rise to an optimal bisabolene titre of 964 mg/L. As significant bisabolene production was detected in the YP control cultivation, a further experiment was conducted to determine whether the organic acids contributed to improvements in titre or whether it was solely due to increased yeast extract. When strain GB2.485 was cultivated at microscale in the optimal media without any organic acids (yeast extract 40 g/L, peptone 10 g/L), the resulting bisabolene titre was 314 ± 25. This was comparable to the titre obtained in the YP control (C3, Figure 1) and around threefold lower than the 964 mg/L predicted for the optimal media by the statistical model (Figure 1B), confirming that bisabolene production is enhanced in cultivations supplemented with organic acids.

#### 3.1.2. Validation of optimal conditions and additional strain testing

To validate the statistical model, the optimal medium conditions were tested at microscale (Figure 2). The levels of those factors, which did not have a significant effect on bisabolene production were set to the lower limit to conserve resources. In addition to strain GB2.485, a further three strains were also grown in the optimised acid medium for comparison (Figure 2). Strain GB2 is another *R. toruloides* strain previously engineered for bisabolene production (Kirby et al., 2021), however, unlike in GB2.485 no mevalonate pathway genes were overexpressed in GB2. Strains EIZS2 and PPS5 were engineered for epi-isozizaene and prespatane production, respectively (Geiselman et al., 2020). For all four strains investigated, biomass accumulation was significantly greater in the control YPD medium compared to the optimised acid medium (Figure 2B). The final OD_600_ values were 38 % lower for strains GB2, EIZS2 and PPS5 cultivated in the optimised medium compared to the control. Interestingly, for strain GB2.485, a lower reduction in biomass accumulation of 18 % was observed (Figure 2). Despite the reduced growth, bisabolene production was significantly greater for optimised media cultivations of strain GB2.485 (*p* = 2.11 × 10^−5^). The final bisabolene titre was 1055 ± 7 mg/L representing a 1.7-fold improvement compared to the YPD control. In addition, this was just 9 % higher than the 964 mg/L predicted by the statistical model (Figure 1C), demonstrating the ability of the derived model to predict performance with a relatively high degree of accuracy. There was no significant difference in bisabolene (*p* = 0.528) and epi-isozizaene (*p* = 0.159) production for strains GB2 and EIZS2, respectively (Figure 2A). Prespatane production by strain PPS5, however, was around 68 % lower in the optimised medium, with a final titre of just 107 mg/L. These results showed that although glucose supplementation was favourable for growth, in three out of four of the engineered *R. toruloides* strains investigated, isoprenoid production was comparable or enhanced when organic acids were used as the main carbon source.

**Figure 2:**
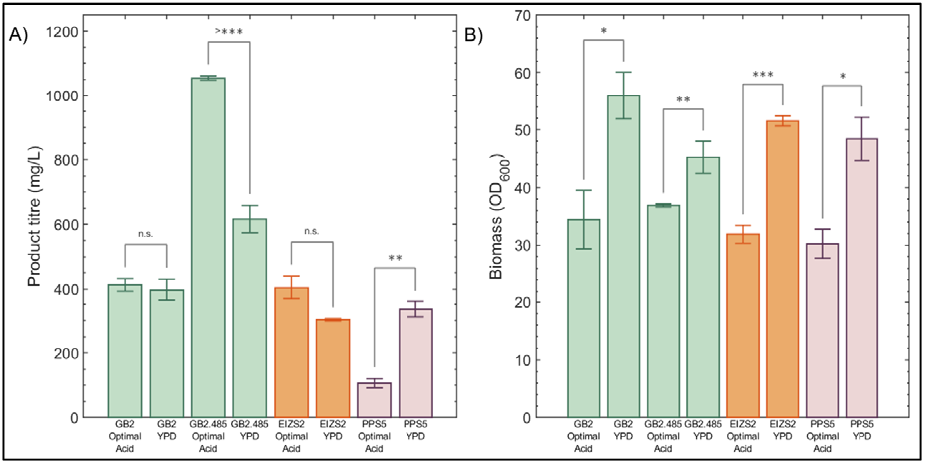
Validation of the statistical model and additional strain testing at microscale. Isoprenoid (A) and biomass (B) accumulation were evaluated at the end of 96 hour cultivations performed using 24-deep-well plates. The isoprenoid products for each strain investigated were as follows: GB2 – bisabolene (green); GB2.485 – bisabolene (green); EISZ2 – epi-isozizaene (orange); PPS5-prespatane (purple). Each strain was cultivated in the optimal acid medium (Yeast extract 40 g/L; Peptone 10 g/L; Succinic acid 9.5 g/L; Acetic acid 5 g/L) and standard YPD (Yeast extract 10 g/L; Peptone 20 g/L; Glucose 20 g/L) as a control. Values are mean ± standard deviation (n = 3; n.s. if *p* > 0.05, * - *p* < 0.05, ** - *p* < 0.01, *** - *p* < 0.001; paired t-test).

### 3.2. Scale up using miniBio 500 bioreactors

#### 3.2.1. Preliminary GB2.485 bioreactor cultivation

The optimal media combination elucidated during the microscale screening was tested at increased scale using a miniBio 500 bioreactor to investigate performance under more industrially relevant conditions as summarised in Figure 3.

**Figure 3:**
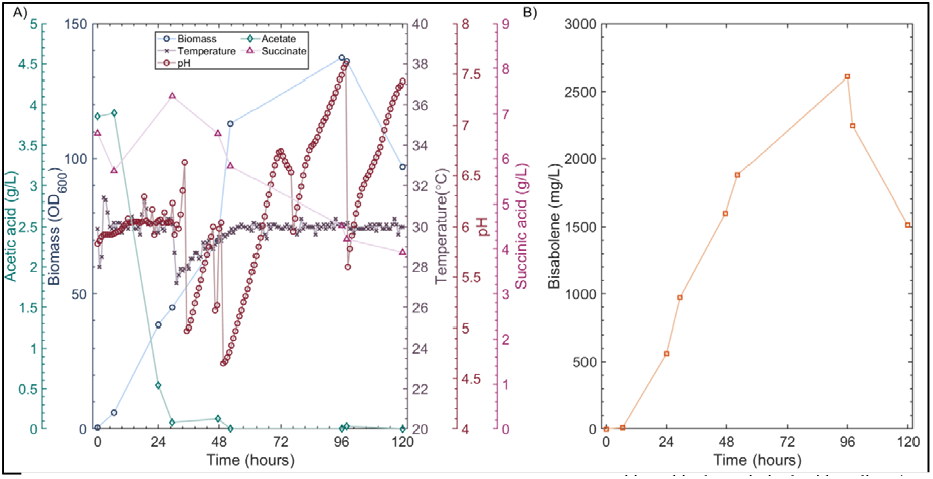
Validation of optimised bioreactor at increased scale. GB2.485 was cultivated in the optimised acid medium (Yeast extract 40 g/L; Peptone 10 g/L; Succinic acid 9.5 g/L; Acetic acid 5 g/L) in a miniBio 500 bioreactor (Applikon Biotechnology). A) Temperature and pH were monitored and controlled online to set points of 30 °C and 6, respectively. Control of pH was achieved through the automatic addition of 2M H_2_SO_4_ or 3M NaOH. Biomass, succinic acid and acetic acid concentrations were monitored via analysis of offline samples taken twice daily. B) Bisabolene production was monitored via GC-MS analysis of dodecane samples taken twice daily.

Despite the application of dissolved oxygen control, oxygen quickly became limiting during exponential growth in the mini-bioreactor. Between 12 and 60 hours, the dissolved oxygen remained well below the setpoint of 30 % (See Supplementary Material). This suggested that a constant air supply throughout cultivations is likely to be beneficial to minimise oxygen limitation. Control of pH was also found to be challenging in the system, with the culture pH varying between 4.5 and 7.5 during the cultivation despite the implementation of pH control with a set point of pH 6. This was partly due to the consumption of the acid substrate causing dramatic increases in culture pH. Back pressure due to the relatively high airflow inside the mini-bioreactor also contributed, through hindering performance of the peristaltic pumps employed for pH control.

Despite process control challenges in the preliminary mini-bioreactor cultivation, a maximum OD_600_ of 137 was achieved at 96 hours, representing a 3.7-fold improvement compared to the 37 ± 0.04 achieved using the same media at microscale (Figure 2B). Bisabolene accumulation was also enhanced 2.5-fold to a maximum titre of 2606 mg/L (Figure 3B).

#### 3.2.2. GB2.485 bioreactor cultivation with pH-controlled acid feeding

Substrate utilisation was readily detectible in the initial bioreactor cultivation, through an increase in culture pH. It was therefore hypothesised that pH controlled organic acid feeding could provide a dual benefit of stabilising conditions inside the reactor and increasing productivity through the provision of additional carbon source, without allowing acids to accumulate to inhibitory levels. To investigate this, a bioreactor cultivation was performed in which a 100 mL, 5X concentrated acid feed (succinic acid, 47 g/L; acetic acid, 25 g/L) was employed for single sided pH control. The results of this experiment are summarised in Figure 4A & B. As increasing the initial yeast extract concentration was found to improve bisabolene production at microscale (Figure 1), it was hypothesised that supplementing the organic acid feed with an additional supply of nutrients may be beneficial for bisabolene production. An additional fed-batch cultivation was therefore performed in which the feed contained all components of the media at 5X concentration (succinic acid, 47 g/L; acetic acid, 25 g/L; yeast extract, 200 g/L; peptone, 50 g/L). The results of this experiment are summarised in Figure 4C & D.

**Figure 4:**
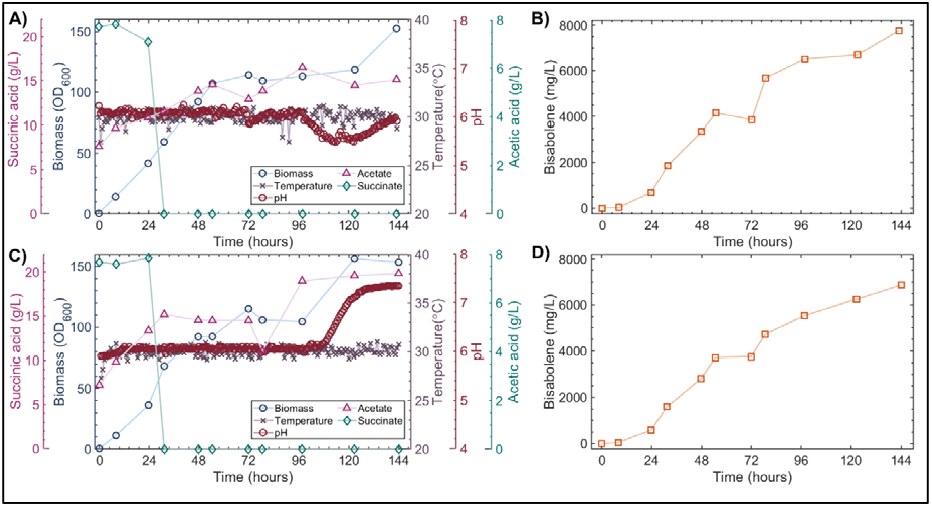
Optimised medium cultivation with pH controlled feeding. Strain GB2.485 was cultivated in the optimised acid medium (Yeast extract 40 g/L; Peptone 10 g/L; Succinic acid 9.5 g/L; Acetic acid 5 g/L) in miniBio 500 bioreactors (Applikon Biotechnology). Single-sided pH control was achieved through automatic addition of either 5X concentrated acid (succinic acid, 47 g/L; acetic acid, 25 g/L; A&B) or 5X concentrated media (succinic acid, 47 g/L; acetic acid, 25 g/L; yeast extract 200 g/L; peptone 50 g/L; C&D), whenever the pH rose above the setpoint of 6. Biomass, succinic acid and acetic acid concentrations were monitored via analysis of offline samples taken twice daily for the concentrated acid (A) and media (C) fed reactors, respectively. Bisabolene production in the concentrated acid (B) and media (D) fed reactors was monitored via GC-MS analysis of dodecane samples taken twice daily.

Prior to the start of the cultivations, the proportional-integral-derivative (PID) settings of the adaptive pH controller of the myControl system (Applikon Biotechnology) were reset, to ensure a slower initial pump rate and reduce the risk of overcorrection. The condenser set up was also modified to prevent pooling of water and thereby ensure unobstructed gas flow through the condenser outlet. These alterations improved pH control in the reactors dramatically, with the pH remaining close to the set-point of 6 for the first 96 hours (Figure 4A &C). In the concentrated acid fed reactor, exponential growth of *R. toruloides* strain GB2.485, was observed between 0 and 54 hours with an OD_600_ of 107 reached after 54 hours of cultivation. The concentration of acetic acid in the cultivation medium rapidly dropped close to 0 g/L during exponential growth and did not increase during substrate feeding. The concentration of succinic acid, on the other hand, increased up to a maximum of 16.5 g/L after 98 hours of cultivation. This indicates that, although succinic acid concentration was deemed to have a significant effect on bisabolene production by the statistical model (Figure 1), it was largely unconsumed at bioreactor scale. The differences in operational mode between scales may have contributed to the discrepancies observed as recent studies have shown that the ability of microscale batch screening experiments to predict performance of larger scale fed-batch processes is limited (Teworte et al., 2022). Future optimisation studies should therefore focus on the incorporation of fed-batch microscale tools to allow closer mimicry of large scale bioreactor conditions from the outset. A plateau in growth was observed between 54 and 123 hours, followed by an increase to an OD_600_ = 153 between 123 and 144 hours (Figure 4A). This increase followed a dip in pH at 120 hours to pH 5.5, as no additional feed was supplied during this period due to the low pH, and no consumption of succinic acid was observed, this additional growth was likely due to consumption of additional carbon sources present within the complex medium.

Bisabolene accumulation was observed throughout the 144 hour cultivation, with a maximum titre of 7758 mg/L observed at 144 hours (Figure 4B). This represents a threefold improvement in titre compared to the batch cultivation with pH control via H_2_SO_4_ and NaOH (Figure 3B). This titre was also around threefold higher that the previously reported maximum microbial bisabolene titre of 2600 mg/L, which was achieved using a hydrolysate containing ∼115 g/L of sugars (Kirby et al., 2021). Whilst succinic acid consumption was limited, acetic acid proved to be an excellent low-cost carbon source for bisabolene production by *R. toruloides* strain GB2.485.

Supplementation of the feed with additional yeast extract and peptone did not further improve growth, with a comparable OD_600_ of 115 observed after 72 hours of cultivation (Figure 4C). As for the concentrated acid fed reactor (Figure 4A), a plateau in growth was observed between 72 and 108 hours. At 108 hours, the concentrated media solution was depleted and the pH rose from the set point of 6 to 7.4. This increase in pH was coupled with an increase in OD_600_ to 156 (Figure 4C), as in the concentrated acid fed cultivation (Figure 4A), succinic acid was not consumed and accumulated in the cultivation medium, this additional growth was therefore likely the result of consumption of additional carbon sources present. Bisabolene production was also comparable in the concentrated media (Figure 4D) and concentrated acid (Figure 4B) fed bioreactors, with final bisabolene titres of 6865 and 7758 mg/L, respectively. As supplementation of the feed with additional yeast extract and peptone did not improve growth and resulted in a slight decrease in bisabolene accumulation, it was deemed inappropriate.

### 3.3. Feasibility of bioconversion of cellulose to organic acids by Ruminococcous flavefaciens

#### 3.3.1. Preliminary Ruminococcous flavefaciens cultivations

Preliminary cultivations of *Ruminococcus flavefaciens* were performed using medium 436 (DSMZ, Germany) under strictly anaerobic conditions. The species was gradually adapted to growth on cellulose as the sole carbon source by replacing glucose with 1 % w/v cellulose. Once metabolism of cellulose could be confirmed through production of a yellow pigment associated with cellulase production (Kopečný and Hodrová, 1997; Sijpesteijn, 1951), the cellobiose was also removed from the medium, yielding cellulose medium. Acid accumulation was evaluated in preliminary cellulose medium cultivations (See Supplementary Material). As the optimal cultivation time was not known, the effect of incubation period was investigated, through measuring total acid concentration after five and seven days. The resulting titres were highly comparable at 3.47 and 3.57 g/L, respectively (See Supplementary Material). Control measurements of the medium without any bacteria were also performed to ensure the acids were produced by the bacterium and not already present in the medium. As only a small amount (∼ 0.1 g/L) of acetic acid and no succinic acid were detected in the control, it can be assumed that the organic acids detected in the cultures were the result of cellulose fermentation by *R. flavefaciens*. Although the total acid concentration was around fourfold lower than the optimal concentration for bisabolene production determined during high-throughput microscale screening (Figure 1), the anaerobic bacteria were found to reliably produce gram per litre quantities of both of the acids of interest from cellulose.

### 3.4. Proof of concept: sequential bioreactor approach for the bioconversion of cellulose to bisabolene

#### 3.4.1. Testing a suitable medium for sequential cultivations

As the concentrations of yeast extract and peptone in the bacterial cellulose medium were 20 and twofold lower, respectively, than those of the optimised acid medium, it was hypothesised that the nutrient content of bacterial medium would be insufficient to support growth of the yeast, especially after fermentation. A bacteria-yeast medium was therefore developed in which the initial concentrations of yeast extract and peptone were increased to 40 and 10 g/L, respectively. *R. flavefaciens* fermentation performance in this bacteria-yeast medium was characterised as summarised in Figure 5.

**Figure 5:**
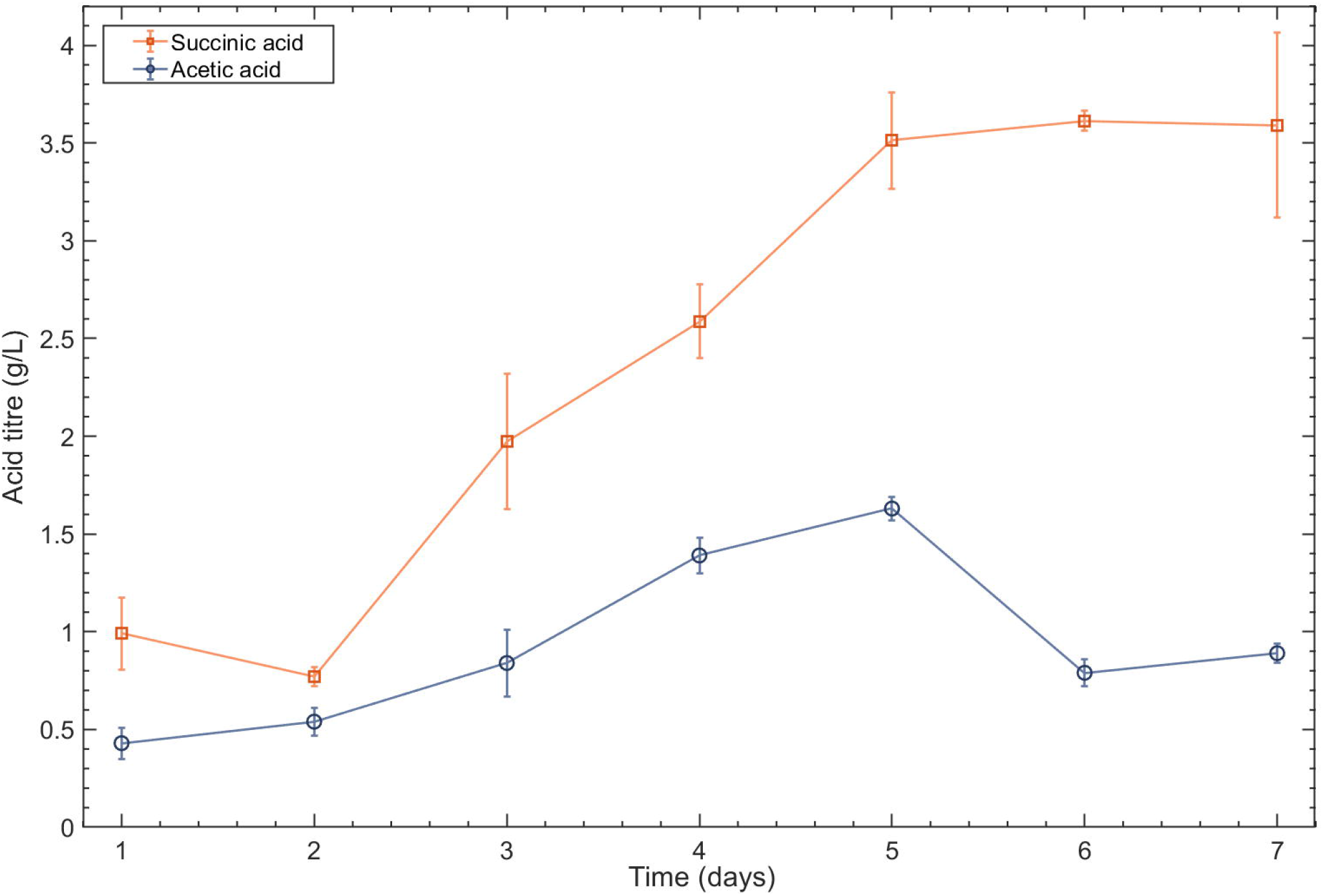

When *R. flavefaciens* was cultivated in the yeast-bacteria medium, succinic acid and acetic acid production was observed during the first five days of cultivation, with maximum titres of 3.51 ± 0.24 and 1.63 ± 0.06 g/L observed after five days, respectively (Figure 5). The succinic acid titre plateaued between five and seven days, and a slight decrease in acetic acid titre was observed. This further confirmed that an incubation time of five days is likely to be optimal for the cellulose fermentation step.

#### 3.4.2. Testing the sequential bioprocess approach

Finally, the sequential bioprocess was tested through first cultivating *R. flavefaciens* in the bacteria-yeast medium for five days. At the end of the five-day fermentations, the resulting cultures were centrifuged and the supernatant used for subsequent microscale cultivations of *R. toruloides* strain GB2.485. The sequential bioprocess is illustrated in Figure 6.

**Figure 6:**
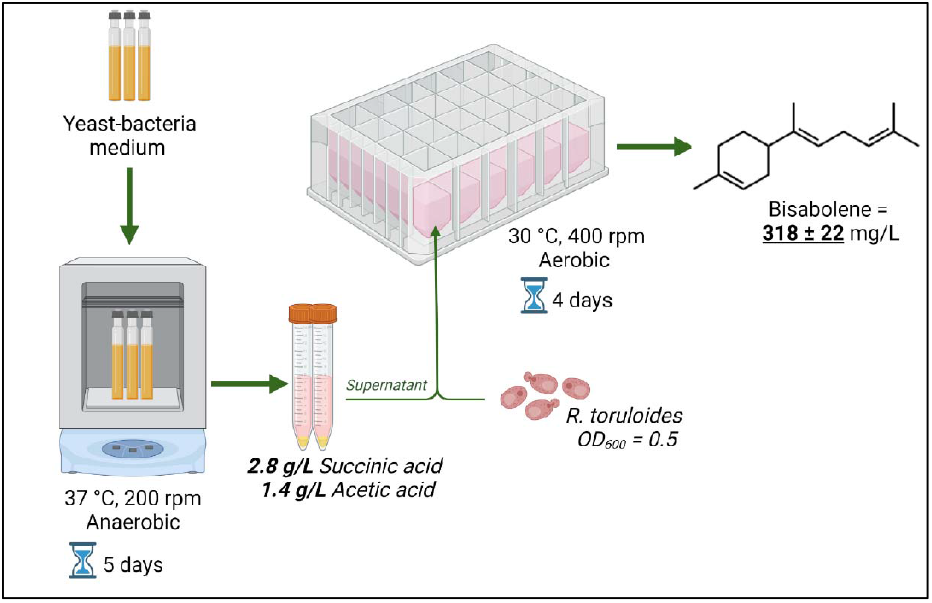
Summary of the sequential bioprocess.

Fermentation of cellulose by *R. flavefaciens* yielded a total organic acid concentration 4.2 g/L at the end of the five day cultivation. Cultivation of *R. toruloides* strain GB2.485 in the fermented bacteria-yeast media resulted in a bisabolene titres of 318 ± 22 mg/L (Figure 6). Although the maximum titre obtained for the fermented bacteria-yeast medium was 3.3- fold lower than the 1055 mg/L achieved using the optimised acid medium at microscale (Figure 2), the product yield coefficients () were highly comparable at 0.076 and 0.073 g/g, respectively. However, it is worth noting that this titre is highly comparable to that of the YP control (C3, Figure 1A) without any organic acids. Future studies using synthetic media will further clarify the potential of this sequential approach using organic acids as sole carbon source. Nevertheless, this early data suggests that through optimisation of the first step to improve the conversion of cellulose and enhance organic acid yields, particularly acetic acid, gram per litre quantities of bisabolene could be generated from cellulose without the need for commercial enzymes to support the growth of our engineered *R. toruloides* strains

In the natural rumen environment, methanogens act as an important hydrogen sink, sequestering hydrogen and CO_2_, yielding methane (Ungerfeld, 2015). This is favourable for overall fermentation performance as reduced co-factors are re-oxidised during methanogenesis allowing fermentation to continue (Ungerfeld, 2015). Previous studies have found that co-cultivation of *Ruminococci* with acetogens, as alternative hydrogen sinks, can improve fermentation performance and shift flux toward acetic acid production (Miller and Wolin, 1995; Williams et al., 1994). Co-cultivation of the cellulolytic rumen bacterium, *Ruminococcus albus*, with an acetogenic bacterium, for example, improved acetic acid accumulation threefold compared to monoculture (Miller and Wolin, 1995). This strategy could potentially improve bisabolene titres in the sequential bioprocess of this study, as bisabolene production was found to occur predominantly during acetic acid metabolism.

## 4. Conclusion

This study demonstrated a proof-of-concept sequential bioprocess for the bioconversion of cellulose into bisabolene. Preliminary microscale screening revealed organic acids as a promising carbon source for bisabolene production by *R. toruloides*, with a maximum titre of 1055 ± 7 mg/L. Implementation of a pH controlled organic acid feeding strategy at bioreactor scale improved titres 7.4-fold to 7758 mg/L, the highest reported microbial titre. *Ruminococcus flavefaciens*, successfully fermented microcrystalline cellulose, yielding 4.2 g/L of organic acids, which were subsequently converted into 318 ± 22 mg/L of bisabolene by *R. toruloides*. Future work should focus on improving organic acid titres and using real lignocellulosic feedstocks.

## 5. Acknowledgements

The authors would like to thank Mr Stuart Martin and Mr Mark Lauchlan at The School of Engineering, University of Edinburgh, UK for their kind assistance and technical support with GC-MS analysis. This work was supported by the Engineering and Physical Sciences Research Council (Grant number EP/R513209/1) and The British Council (Grant Number: 527429894). The *R. toruloides* strains used in this work were developed by the Joint BioEnergy Institute through the U.S. Department of Energy, Office of Science, Office of Biological and Environmental Research, through contract DE-AC02-05CH11231 between Lawrence Berkeley National Laboratory and the U.S. Department of Energy.

